# CSF1R inhibition by PLX5622 reduces pulmonary fungal infection by depleting MHCII^hi^ interstitial lung macrophages

**DOI:** 10.1101/2022.11.18.517077

**Authors:** Sally H. Mohamed, Eliane Vanhoffelen, Man Shun Fu, Emilie Cosway, Graham Anderson, Greetje Vande Velde, Rebecca A. Drummond

## Abstract

PLX5622 is a small molecular inhibitor of the CSF1 receptor (CSF1R) and is widely used to deplete macrophages within the central nervous system (CNS). However, recent reports have indicated that PLX5622 may affect myeloid cells in other organs including the bone marrow and spleen. We investigated the impact of PLX5622 treatment in wild-type C57BL/6 mice and discovered that one-week treatment with PLX5622 was sufficient to deplete interstitial macrophages in the lung and brain-infiltrating Ly6C^low^ patrolling monocytes, in addition to CNS-resident macrophages. These cell types were previously indicated to act as infection reservoirs for the pathogenic fungus *Cryptococcus neoformans*. We therefore took advantage of PLX5622-mediated depletion of these myeloid cell subsets to examine their functional role in *C. neoformans* lung infection and extrapulmonary dissemination. We found that PLX5622-treated mice had significantly reduced fungal lung infection and reduced extrapulmonary dissemination to the CNS but not to the spleen or liver. Fungal lung infection mapped to MHCII^hi^ interstitial lung macrophages, which underwent significant expansion during infection following monocyte replenishment and not local division. Although PLX5622 depleted CNS infiltrating patrolling monocytes, these cells did not accumulate in the fungal-infected CNS following pulmonary infection. In addition, Nr4a1-deficient mice, which lack patrolling monocytes, had similar control and dissemination of *C. neoformans* infection to wild-type controls. Our data demonstrate that PLX5622 may have a beneficial effect in the control of intracellular replicating pathogenic fungi that utilise CSF1R-dependent myeloid cells as infection reservoirs.

## Introduction

Myeloid cell population size is predominantly controlled by the growth factor CSF1 (M-CSF) [1]. Expression of the CSF1 receptor (CSF1R) broadly identifies the myeloid lineage, but is particularly highly expressed by neutrophils, monocytes and macrophages [2]. Different subsets of macrophages and monocytes exhibit varying degrees of sensitivity to CSF1, which appears to correlate with the dependency on CSF1R for survival [2]. For example, maintenance and development of central nervous system (CNS)-resident microglia is highly dependent on the CSF1R, since mutation or deletion of this receptor results in complete ablation of microglia numbers in rodents and humans [3–5]. In the lung, alveolar and interstitial macrophages express similar levels of CSF1R but have divergent uptake rates of CSF1, with interstitial macrophages more readily responsive to CSF1 stimulation [2]. However, genetic deletion of CSF1R in mice and rats does not reduce total numbers of lung macrophages [3, 5].

Since CNS-resident macrophages are highly dependent on CSF1R signalling for survival, chemical inhibitors of CSF1R are an effective tool to deplete these macrophage populations [6]. The most specific CSF1R inhibitor is PLX5622, which has greater brain penetrance than other CSF1R inhibitors (e.g. PLX3397) [7, 8]. When given orally, PLX5622 rapidly depletes CNS-resident macrophages (including parenchymal microglia and border macrophage populations) [8]. Several studies have used PLX5622-treatment models to demonstrate the critical role of microglia in controlling CNS infections and neuroinflammation [8–10]. However, controversy over the specificity of PLX5622 has emerged in recent years [11, 12]. One study showed that PLX5622 caused significant alterations to both myeloid and lymphoid populations within the spleen and bone marrow, and macrophages derived from the bone marrow of PLX5622-treated mice had impaired production of IL-1β [11]. Many of these effects remained 3 weeks after PLX5622 treatment had stopped, indicating at long-term consequences of disrupting CSF1R-dependent myeloid compartments [11]. In other studies, monocyte numbers were reduced by PLX5622 treatment although these effects may be subset and context specific [10, 13, 14], since other studies have found no effect on either monocyte or non-CNS macrophage numbers [15, 16].

Tissue-resident macrophages and monocytes are integral to the pathogenesis of the fungal infection caused by *Cryptococcus neoformans* [17]. This opportunistic fungal pathogen causes lung infections and life-threatening meningitis in patients with compromised T-cell immunity. Cryptococcal meningitis causes 19% of all AIDS-related deaths globally [18]. Macrophages and monocytes have a poor capacity to kill *C. neoformans* without the help of IFNγ-producing CD4 T-cells. In the absence of Th1 immunity, monocytes and macrophages are susceptible to intracellular infection, marked by strong upregulation of arginase and an ‘M2’ like functional phenotype [17]. *C. neoformans*-infected monocytes are thought to contribute towards dissemination of the infection, especially to the brain and CNS [19, 20]. Tissue-resident macrophages are also an important reservoir for *C. neoformans* in both the brain and lung. We recently found that targeted microglia depletion could reduce fungal brain infection, since microglia shielded *C. neoformans* from copper starvation within the CNS [21]. In the lung, *C. neoformans* produces a secreted peptide called CPL-1 that drives arginase expression within interstitial macrophages to promote intracellular growth [22]. The ability of this fungus to access and infect monocytes and macrophages is therefore critical for the infection lifecycle.

In the present work, we characterised the impact of PLX5622 treatment on numbers of myeloid cells in the CNS, spleen, lung, blood and bone marrow. We discovered that while PLX5622 primarily depleted CNS-resident macrophages, there was additional specific depletion of CNS-infiltrating patrolling monocytes and lung interstitial macrophages. Since these myeloid populations were previously implicated in the pathogenesis of *C. neoformans* infection, we investigated whether PLX5622 could have a therapeutic effect for this fungal infection and explored the functional roles of these cells during lung infection and extrapulmonary dissemination.

## Results

### Interstitial lung macrophages are depleted by PLX5622

PLX5622 is used to deplete tissue-resident macrophages within the CNS that are highly dependent on CSF1R signalling for survival [6]. However, some reports have suggested that PLX5622 treatment has off-target effects, particularly within the spleen, bone marrow and lung [11]. To independently verify those results, we measured the number of macrophages in brains, meninges, spleens, bone marrow and lungs of mice given control diet or high concentration (1200 ppm) PLX5622 diet for 1 week, using flow cytometry (Fig S1). We found that PLX5622-treated mice had more than 90% loss of microglia and other CNS-resident macrophages (Fig 1A), as previously described. In addition, we also found a significant loss of splenic macrophages in the PLX5622-treated animals (Fig 1B), but no effect on macrophages within the bone marrow (Fig 1C). In the lung, we found that PLX5622 specifically depleted interstitial macrophages whereas alveolar macrophage numbers were unaffected (Fig 1D). Taken together, our data show that high dose PLX5622 has the most profound effect on CNS-resident macrophages but also depletes splenic and interstitial lung macrophages.

**Figure 1:**
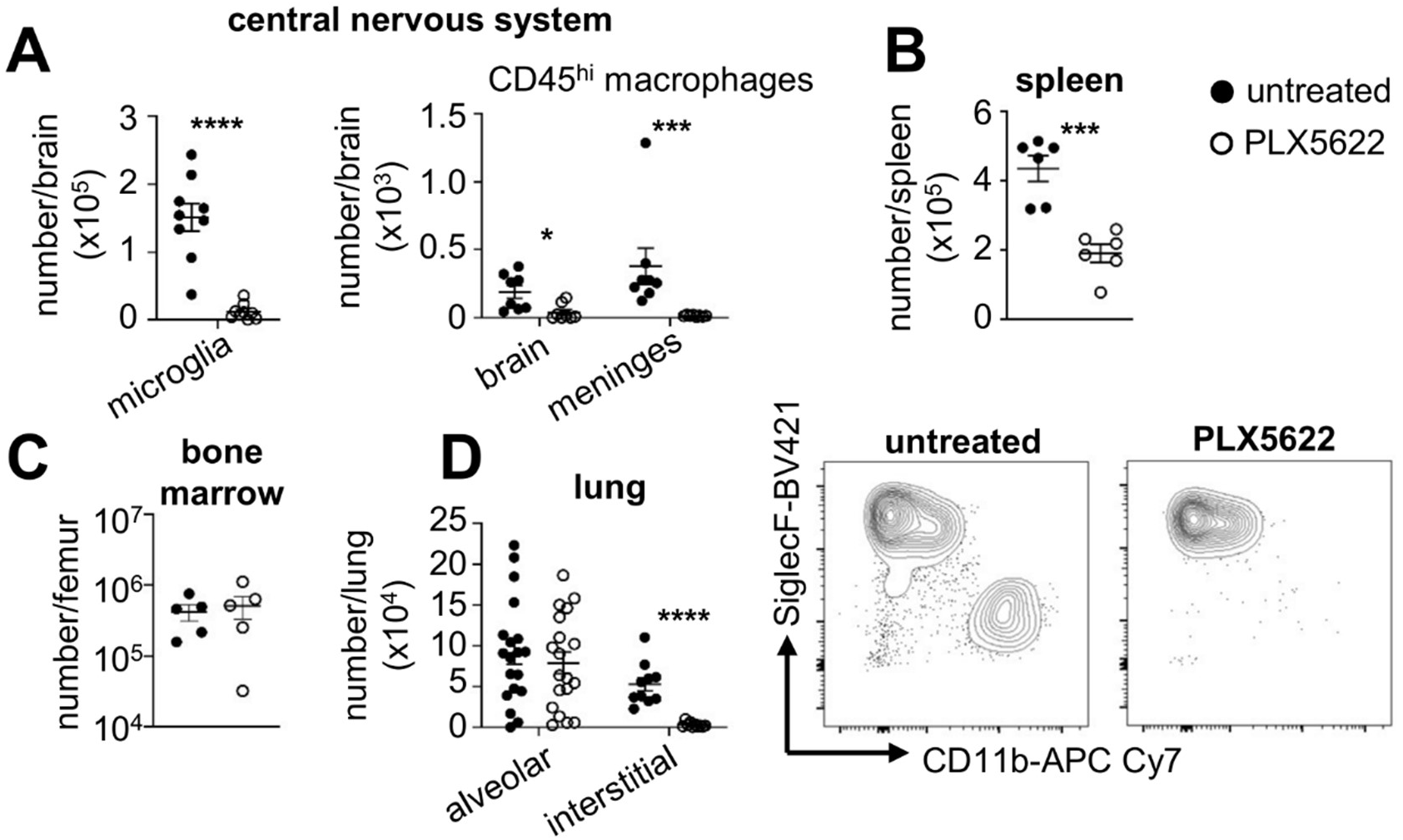
PLX5622 depletes macrophages within the CNS, spleen and lung. Wild-type C57BL/6 mice were given PLX5622 diet (1200ppm) or control diet for 7 days prior to measurement of macrophage numbers in (A) brain and meninges (microglia n=9 per group; macrophages n=8 per group), (B) spleen (n=6 mice per group) and (C) lung (alveolar n=19 per group; interstitial n=10 per group) by flow cytometry. Example lung plots are gated on live CD45^+^CD64^+^MerTK^+^ singlets. Gating strategies for each macrophage population is shown in Fig S1. Data is pooled from 2-4 independent experiments and analysed by unpaired t-tests. ***P<0.005, ****P<0.0001.

### PLX5622 depletes patrolling monocytes in an organ-specific manner

Monocytes are a heterogeneous group of circulating myeloid cells. The two major functional classes of these are classical inflammatory monocytes (Ly6C^hi^ in mice) or non-classical patrolling monocytes (Ly6C^low^) [23]. Some studies have indicated that PLX5622 induces depletion of both or one of these monocyte subsets [10, 13, 14], while others have observed no effect on either subset [15, 16]. To determine whether PLX5622 causes loss of monocytes at the steady-state, we quantified the number of Ly6C^hi^ classical monocytes and Ly6C^low^ patrolling monocytes in the blood, lung, spleen, bone marrow and CNS after 1 week of PLX5622 treatment. We found that PLX5622 had no effect on the number of circulating monocytes in the blood, lung, spleen or bone marrow (Fig 2A-D). In contrast, we found a significant depletion of Ly6C^low^ monocytes in the brain of PLX5622-treated mice (Fig 2E), but no effect in the meninges (Fig 2F). Classical Ly6C^hi^ monocytes remained similar to untreated controls in all tissues examined (Fig 2). Our data show that CNS-infiltrating Ly6C^low^ patrolling monocytes are depleted by PLX5622, which may be secondary to microglia depletion as circulating monocytes were unaffected.

**Figure 2:**
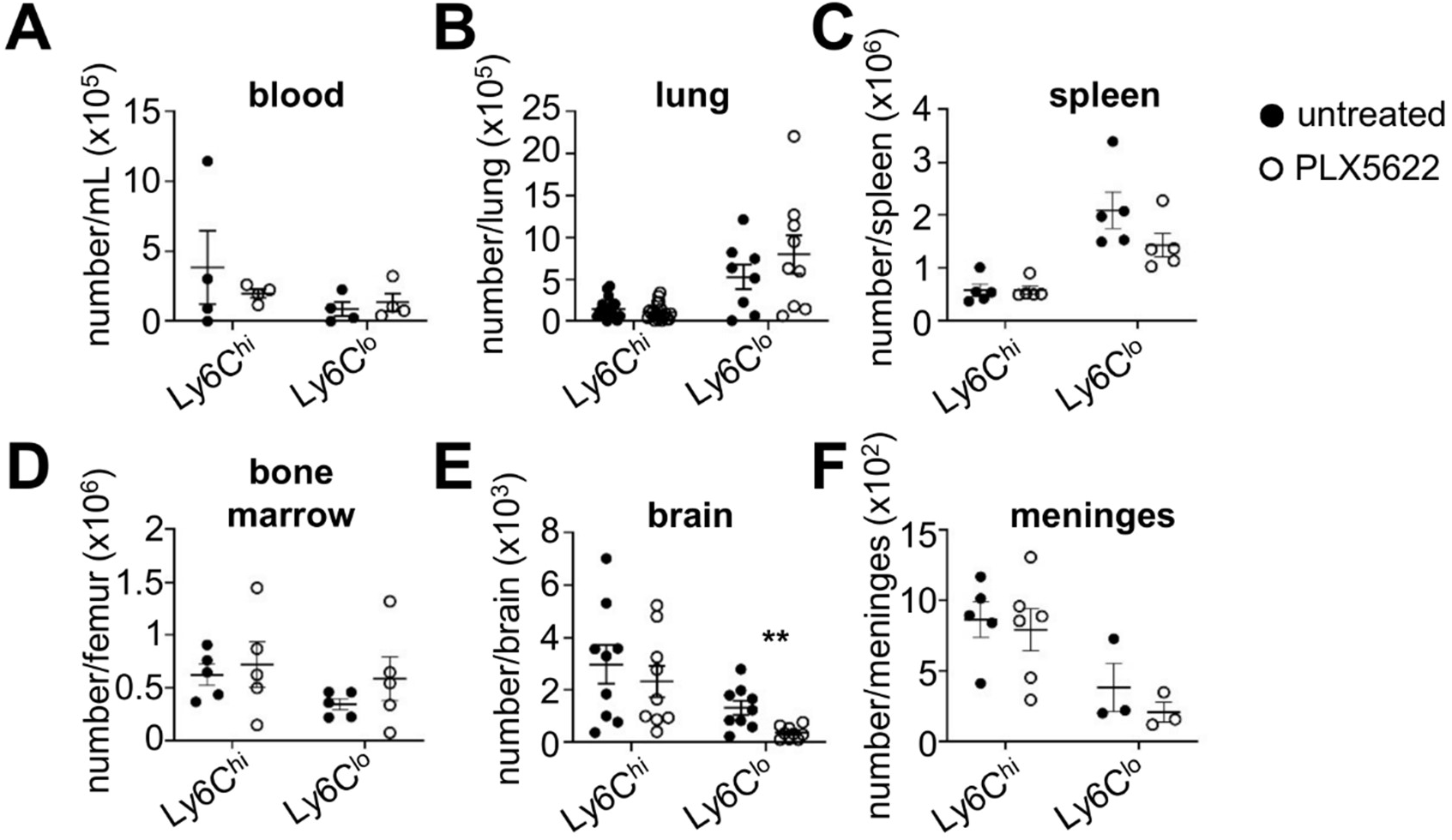
CNS-infiltrating Ly6C^low^ monocytes are depleted in PLX5622-treated mice. Wild-type C57BL/6 mice were given PLX5622 diet (1200ppm) or control diet for 7 days prior to measurement of monocyte numbers in (A) blood (n=4 per group), (B) lung (Ly6C^hi^ monocytes n=19 per group; Ly6C^low^ monocytes n=9 per group), (C) brain (n=9 per group) by flow cytometry. Gating strategies for each monocyte population is shown in Fig S1. Data is pooled from 2-4 independent experiments and analysed by unpaired t-tests.

### PLX5622 reduces fungal lung infection

As we had found that PLX5622 specifically depleted interstitial macrophages in the lung, we took advantage of this to examine the functional role of these cells in *C. neoformans* lung infection, in which interstitial macrophages were recently identified as an infection reservoir [22]. We treated wild-type mice with PLX5622 for 1 week and then performed an intranasal infection with *C. neoformans* prior to measuring fungal burdens in the lung at day 21 post-infection (Fig 3A). We found that PLX5622-treatment led to a significant reduction in lung fungal burden (Fig 3B) and to a non-significant reduction in lung lesion volume (Fig S2), but had no effect on the recruitment of inflammatory cells to the infected lung including monocytes, neutrophils and eosinophils (Fig 3C). We also found no difference in the numbers of lymphocytes, either at steady-state or following infection (Fig 3D). Lymphocyte numbers were also unaffected in the blood, bone marrow and spleen of PLX5622-treated mice (Fig S3). PLX5622-induced depletion of interstitial macrophages was maintained up to day 21 post-infection (Fig 3C) with no effect observed on alveolar macrophages, both in number (Fig 3C) and their expression of activation markers MHCII and F480 (Fig S4). PLX5622 also did not directly have an antifungal effect, as yeast grown in PLX5622 (at concentrations previously measured in brain and serum of treated mice [24]) grew similarly to untreated yeast cultures (Fig S5). Taken together, this data shows that PLX5622 treatment reduces fungal lung infection, which is correlated with a specific loss of interstitial macrophages.

**Figure 3:**
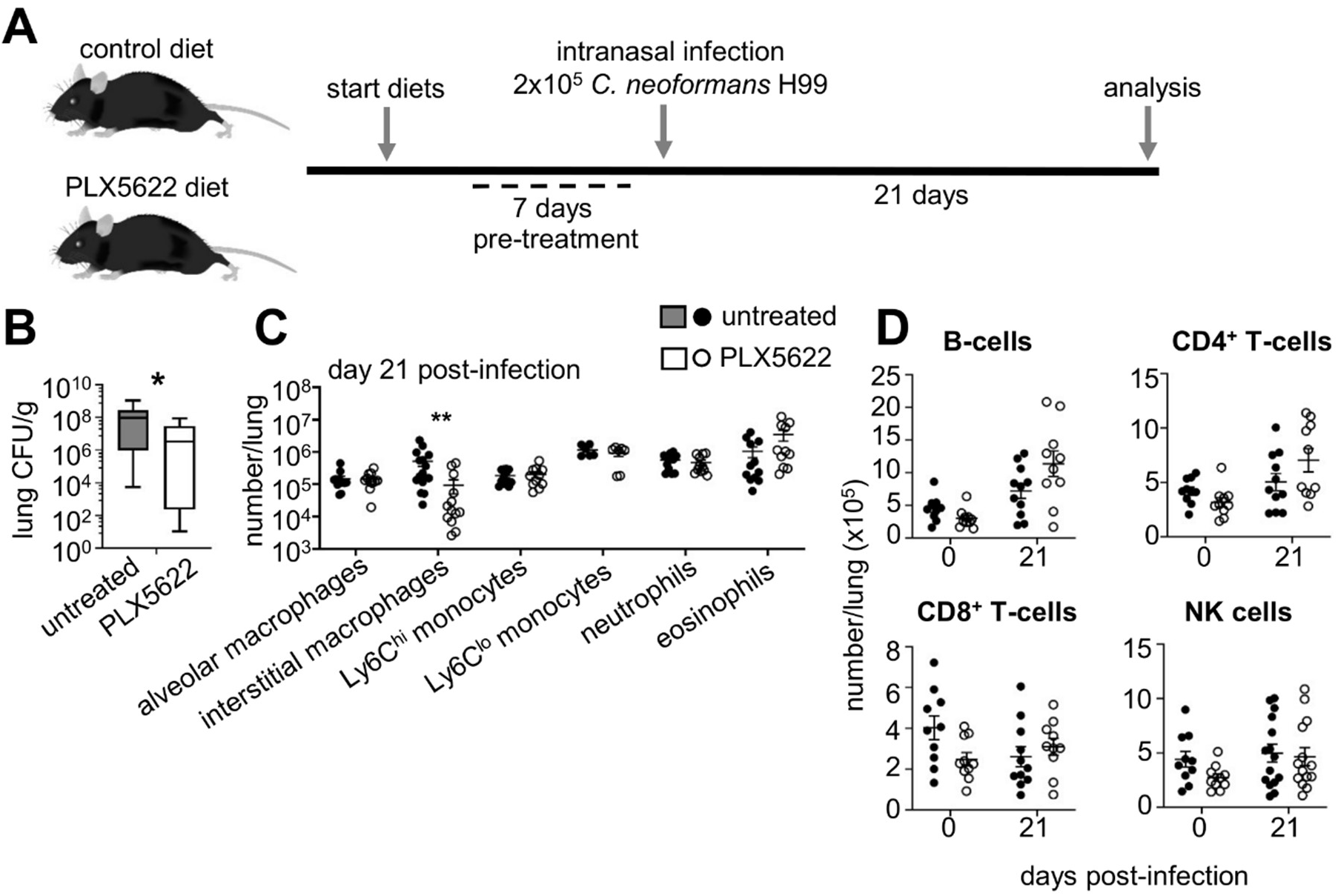
PLX5622 treatment reduces *C. neoformans* lung infection. (A) Schematic of PLX5622 treatment and infection model. Wild-type C57BL/6 mice were pre-treated for 7 days with PLX5622 diet or control diet, then infected intranasally with *C. neoformans*. Mice were analysed up to 21 days post-infection, control and PLX5622 diets were maintained throughout. (B) Lung fungal burdens at day 21 post-infection in untreated (n=10) and PLX5622-treated (n=10) mice. Data pooled from 2 independent experiments and analysed by Mann Whitney U-test, *P<0.05. (C) Flow cytometry was used to determine number of indicated immune cells in the lung at day 21 post-infection (n=12-16 per group). Data is pooled from 3-4 independent experiments and analysed by two-way ANOVA, **P<0.01. (D) Flow cytometry was used to determine the number of indicated lymphocyte populations in the lungs of uninfected (n=10 per group) and at day 21 post-infection (n=10-11 per group) of untreated and PLX5622-treated mice. Data is pooled from 2-3 independent experiments.

### Extrapulmonary dissemination to the CNS is reduced by PLX5622 treatment

Previous work has shown that the rate of lung infection can determine the kinetics of extrapulmonary dissemination to the CNS [25]. Since we had observed that PLX5622 reduced fungal lung infection (Fig 3B), we measured fungal burdens in other tissues to examine the effect of PLX5622 treatment on fungal dissemination from the lung. After day 21 post-intranasal infection, we found that fungal burdens in the brain were significantly reduced in PLX5622-treated animals compared to untreated controls (Fig 4A). In contrast, extrapulmonary dissemination to the spleen and liver were unaffected by PLX5622 treatment (Fig 4A). In the brain, PLX5622-treated mice had significantly reduced numbers of microglia and macrophages (Fig 4B) similar to uninfected mice (Fig 1A). Numbers of recruited inflammatory cells (neutrophils, inflammatory monocytes) were unaffected by PLX5622 treatment (Fig 4B). Numbers of patrolling Ly6C^low^ monocytes were reduced in the infected brains of PLX5622-treated mice (Fig 4B), similar to what we observed in uninfected mice.

**Figure 4:**
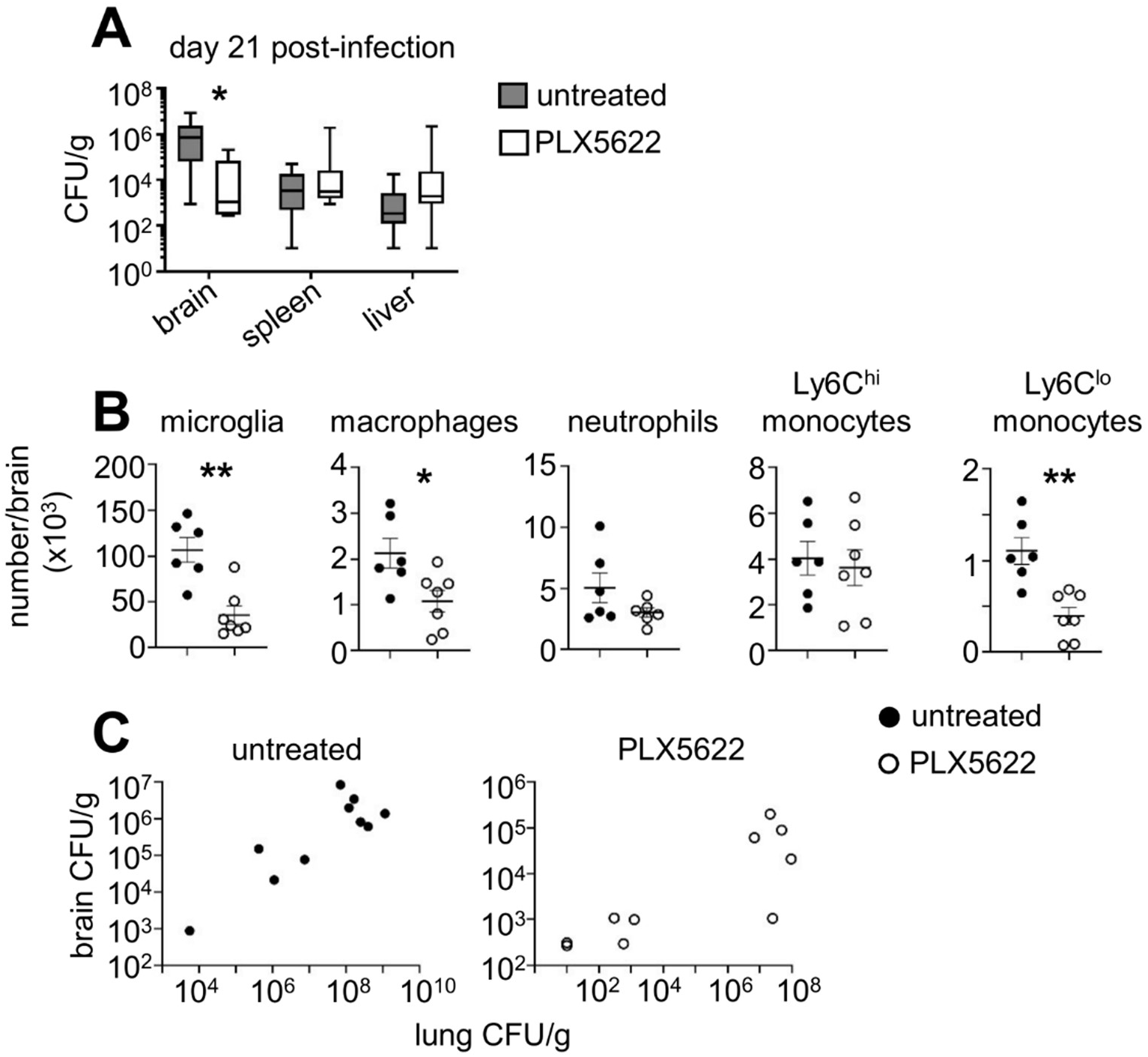
PLX5622 treatment reduces fungal brain infection. (A) Fungal burdens in the indicated organs at day 21 post-infection in untreated and PLX5622-treated mice (brain n=10 mice per group; liver/spleen n=7 mice per group). Data pooled from 2-3 independent experiments and analysed by Mann Whitney U-tests. *P<0.05. (B) Total numbers of indicated myeloid cells in the brain of untreated (n=6) and PLX5622-treated (n=7) mice at day 21 post-infection. Data is pooled from 2 independent experiments and analysed by unpaired t-tests. *P<0.05, **P<0.01. (C) Correlation between fungal burdens in lung and brain of same mice that were either untreated (n=10) or PLX5622-treated (n=10) at day 21 post-infection. Data pooled from 2 independent experiments.

We recently showed that microglia support *C. neoformans* brain infection by acting as a fungal reservoir during acute meningitis (intravenous infection) [21]. Loss of microglia in the PLX5622-treated mice may therefore cause a reduction in brain fungal burdens by removing this growth niche, in addition to reduced extrapulmonary dissemination from the lung, or the loss of microglia independently reduces brain fungal burden (as we previously observed in the intravenous infection model). To explore these possibilities, we first correlated lung and brain fungal burdens in the same mice to determine whether higher lung burdens predicted greater brain infection (and thus increased extrapulmonary dissemination). Indeed, we found that mice with higher lung infection had greater brain infection, and PLX5622 treatment did not disrupt this relationship (Fig 4C). These data indicate that PLX5622-treated mice had reduced brain infection due to reduced lung burdens, although we cannot rule out an additional independent effect of microglia depletion on control of brain infection. Taken together, these data show PLX5622 treatment causes a reduction in extrapulmonary dissemination to the CNS but not to the spleen or liver.

### *C. neoformans* preferentially infects MHCII^hi^ lung interstitial macrophages

Since PLX5622 specifically depleted interstitial macrophages in the lung and this correlated with a reduction in lung infection and dissemination to the CNS, we further characterised these cells in response to *C. neoformans* infection. Lung interstitial macrophages comprise a heterogeneous population of different functional subsets, some which associate with nerve bundles and others with blood vessels [26, 27]. We used MHC Class II to subdivide interstitial macrophages into two subsets (Fig 5A) and analysed their responses to fungal infection. There was a greater number and ratio of MHCII^hi^ interstitial macrophages in the lung, both at the steady-state and during fungal infection (Fig 5B). We found that both subsets of interstitial macrophages expanded at day 21 post-*C. neoformans* infection, although this only achieved statistical significance for the MHCII^hi^ subset (Fig 5B). PLX5622 treatment significantly ablated the MHCII^hi^ interstitial macrophages, and had less of an impact on the less numerous MHCII^low^ interstitial macrophages (Fig 5C).

**Figure 5:**
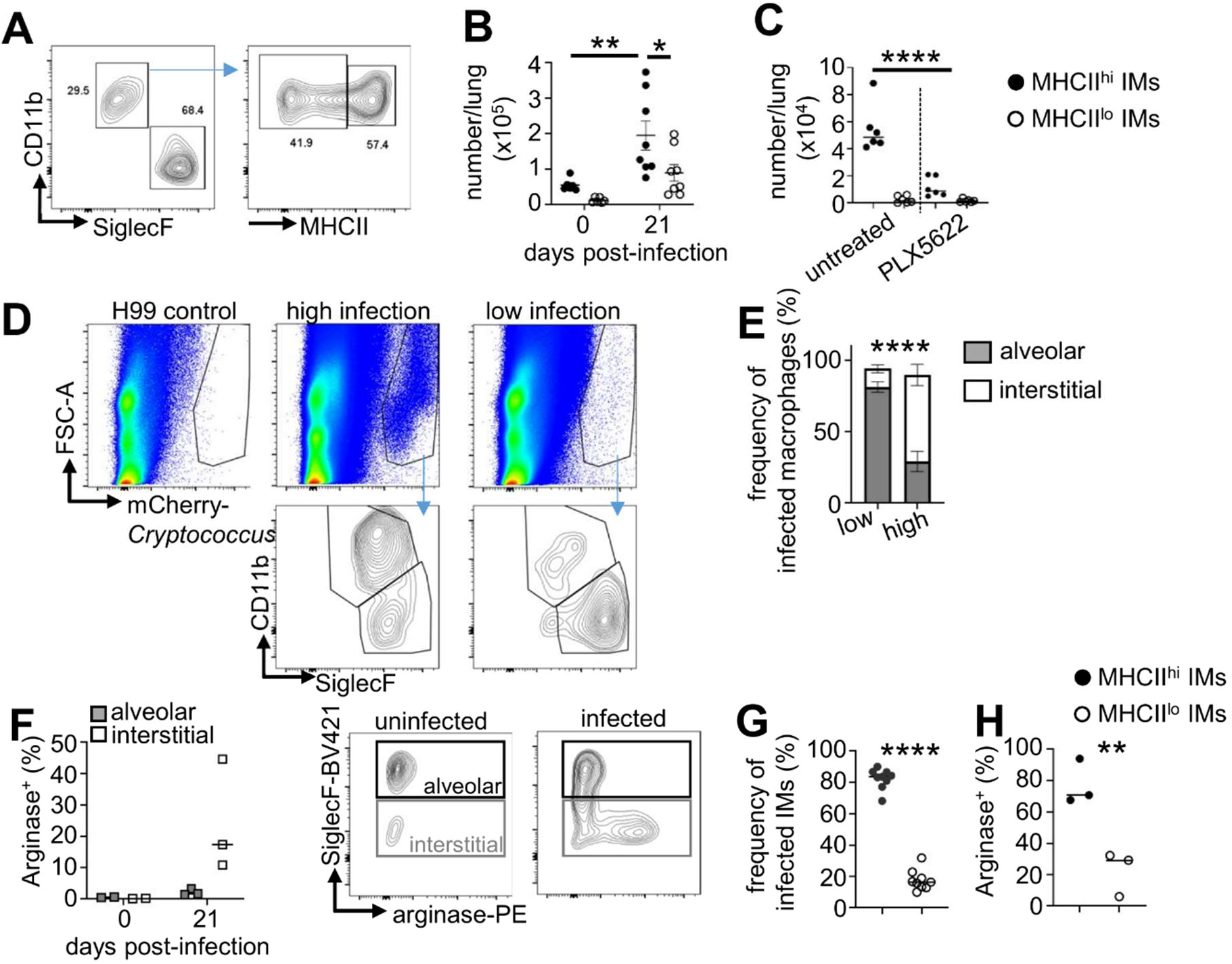
MHCII^hi^ interstitial lung macrophages are susceptible to *C. neoformans* infection. (A) Example flow cytometry of interstitial macrophage gating strategy used in this study. (B) Total number of MHCII^hi^ and MHCII^low^ interstitial macrophages in the lung of uninfected (n=6 per group) and infected (n=8 per group) mice. Data pooled from 2 independent experiments and analysed by two-way ANOVA. *P<0.05, **P<0.01. (C) Total number of interstitial macrophage subsets in the lung of uninfected mice that were either untreated (n=6 mice) or PLX5622-treated (n=6 mice). Data pooled from 2 independent experiments and analysed by one-way ANOVA. ****P<0.0001. (D) Example plots showing ungated lung samples from mice infected with non-fluorescent *C. neoformans* (H99; gating control), and mice infected with mCherry-*C. neoformans* that were grouped as having ‘high infection’ or ‘low infection’. Yeast were further gated as CD45^+^CD64^+^MerTK^+^ to isolate infected macrophages, which were further split into alveolar and interstitial macrophage populations (bottom row). (E) Frequency of infected alveolar and interstitial macrophages (within total infected macrophage gate, CD45^+^CD64^+^MerTK^+^) in mice (n=9 total) with high (n=3) or low (n=6) infection of the lung. Data pooled from 3 independent experiments and analysed by two-way ANOVA. ****P<0.0001. (F) Frequency of arginase expression within alveolar and interstitial macrophages in uninfected (n=2 mice per group) and infected (n=3 mice per group). Example plots are gated on CD64^+^MerTK^+^ macrophages. Data is from a single experiment. (G) Frequency of *C. neoformans* infection within MHCII^hi^ and MHCII^low^ interstitial macrophages (within total infected interstitial macrophages). Data pooled from 3 independent experiments (n=9 total mice), and analysed by unpaired t-test. ****P<0.0001. (H) Frequency of arginase expression in MHCII^hi^ and MHCII^low^ interstitial macrophages in the *C. neoformans* infected lung. Data is from a single experiment (n=3 mice per group).

We next examined the interaction of interstitial macrophages with fungi following infection, using mCherry-expressing *C. neoformans* to track its localisation to different myeloid cell populations in the lung after intranasal infection (Fig 5D). These experiments revealed that *C. neoformans* primarily localised within interstitial macrophages compared to alveolar macrophages when the frequency of detected yeast cells was higher (>0.05% total lung events) (Fig 5E). In contrast, mice with a lower fungal lung burden (<0.05% of total lung events) had a greater proportion of uptake by alveolar macrophages compared to interstitial macrophages (Fig 5E). These data indicate that interstitial macrophages may be more susceptible to intracellular infection, which promotes a greater level of infection. To test that, we measured expression of arginase using intracellular flow cytometry, since arginase-expressing macrophages have been previously shown to support intracellular fungal growth and have poor killing capacity. Indeed, we found that arginase expression was dramatically increased in interstitial macrophages following infection compared to alveolar macrophages (Fig 5F), supporting recent work that also found interstitial macrophages were primarily susceptible to intracellular fungal infection via arginase induction [22]. Next, we examined which interstitial macrophage subset had greater uptake/association with *C. neoformans*. We found that MHCII^hi^ interstitial macrophages were the predominant subset interacting with fungi, regardless of the total frequency of infected cells (Fig 5G). In line with that, MHCII^hi^ interstitial macrophages expressed higher levels of arginase than MHCII^low^ interstitial macrophages (Fig 5G). Taken together, our data show that MHCII^hi^ interstitial macrophages undergo a significant expansion following *C. neoformans* infection and that these cells are susceptible to intracellular infection.

### Interstitial macrophage expansion is driven by monocytes and not local proliferation

As we had observed increases in the size of the interstitial macrophage population in response to fungal infection, we sought to understand the main driver of this expansion. Type-2 cytokines such as IL-4 can promote macrophage proliferation [28] and type-2 immune responses are a key feature of *C. neoformans* lung infections. Indeed, we also observed increases in the concentration of IL-4 and IL-13 in the lung following infection (Fig S6). However, *C. neoformans* intracellular infection can also introduce cell cycle blocks in macrophages [29]. We therefore measured interstitial macrophage proliferation in the uninfected and infected lung using Ki67 as a marker for dividing cells. In uninfected mice, we found little proliferation occurring in alveolar macrophages and eosinophils, with a greater proportion of interstitial macrophages staining positive for Ki67 at baseline (Fig 6A). Upon infection, we found that this level significantly decreased, for both eosinophils and interstitial macrophages (Fig 6A). The level of proliferation did not significantly differ between MHCII^low^ and MHCII^hi^ interstitial macrophage subsets (Fig S7). Therefore, the expansion of interstitial macrophages is not driven by proliferation, and instead fungal infection decreases the baseline proliferation of these cells. In line with that, we found no difference in the type-2 response between untreated and PLX5622-treated lungs, measured either by total IL-4/13 in lung homogenates (Fig S6) or by eosinophil recruitment (Fig 3C).

**Figure 6:**
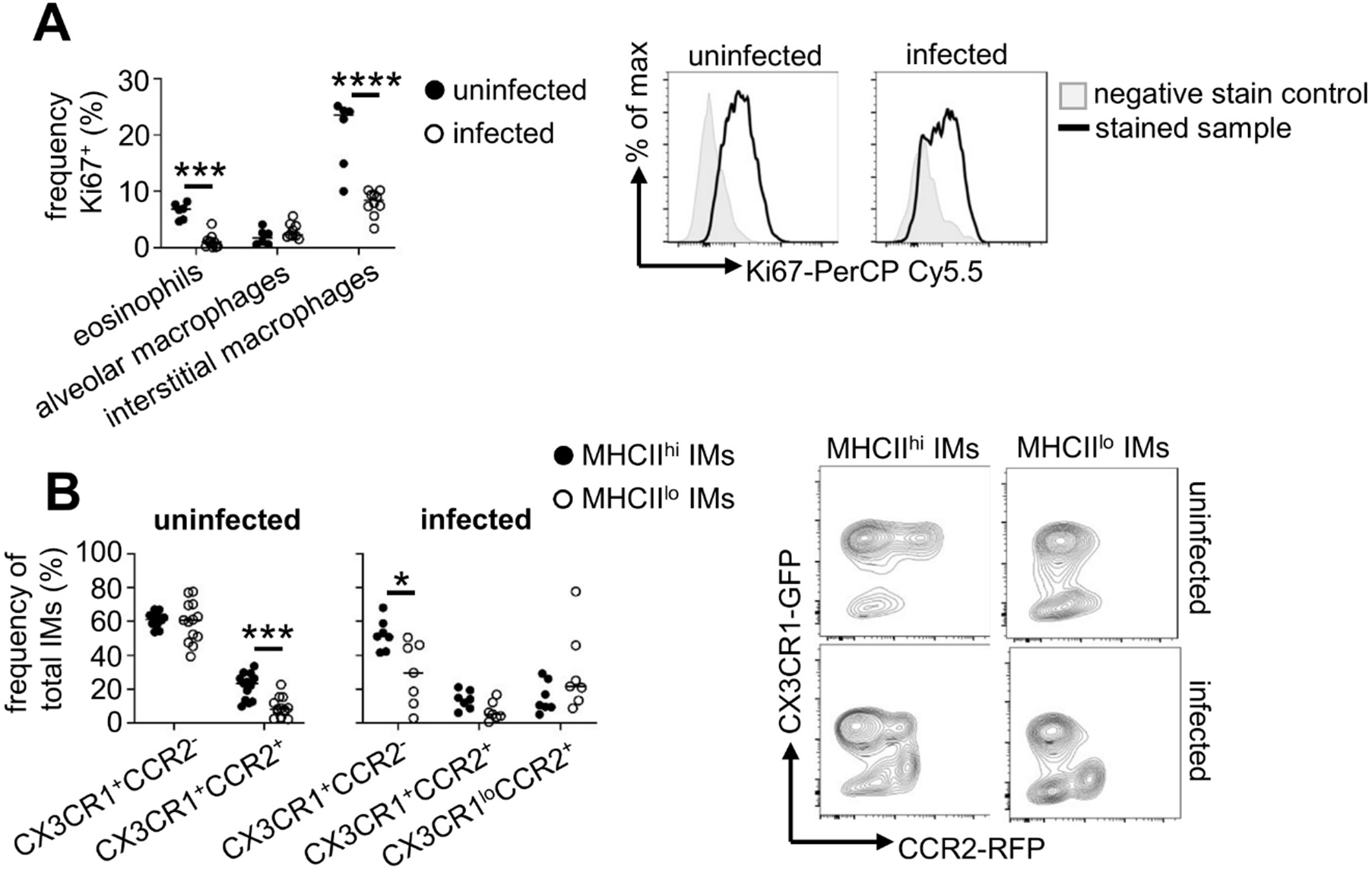
Interstitial macrophage expansion during *C. neoformans* infection is due to monocyte replenishment and not local division. (A) Frequency of proliferating (Ki67+) eosinophils, alveolar and interstitial macrophages in the lungs of uninfected (n=6) and at day 21 post-infection (n=9) mice. Example histograms are gated on interstitial macrophages for infected lung stained with Ki67 (black line) or the fluorescence minus one control (grey filled histogram). Data is pooled from 2 independent experiments and analysed by two-way ANOVA. ***P<0.005, ****P<0.0001. (B) Frequency of interstitial macrophages positive for CX3CR1 and CCR2 expression in the uninfected (n=13 mice) and day 21 post-infected lung (n=7 mice). Example plots are gated on the indicated interstitial macrophage sub-populations. Data is pooled from 2-3 independent experiments and analysed by two-way ANOVA. *P<0.05, ***P<0.005.

Since proliferation was not the main driver of interstitial macrophage expansion during fungal infection, we instead examined whether monocytes differentiating into interstitial macrophages were the primary cause of interstitial macrophage expansion. For that, we used *Cx3cr1-GFP-Ccr2-RFP* mice to track expression of tissue-resident (CX3CR1) and monocyte (CCR2) markers in interstitial macrophages during fungal lung infection. At baseline, MHCII^hi^ interstitial macrophages were mostly CX3CR1^+^CCR2^-^ (tissue-resident) with a smaller population of CX3CR1^+^CCR2^+^ monocyte-derived cells (Fig 6B). In contrast, MHCII^low^ interstitial macrophages lacked a CCR2^+^ population, indicating that monocytes are not a significant contributor to this population (Fig 6B). We did not detect CX3CR1 or CCR2 expression in alveolar macrophages or eosinophils (Fig S8). After fungal infection, we found a population of CX3CR1^low^CCR2^+^ cells emerged in both interstitial macrophage subsets (Fig 6B), which likely represents immature monocyte-derived cells that contribute to the interstitial macrophage pool in the lung following infection.

Taken together, our data show that the interstitial macrophage population dynamically changes in response to fungal infection, undergoing monocyte-driven expansions that may be driven by reduced capacity to proliferate as a result of intracellular infection.

### Patrolling monocytes play a minor role in extrapulmonary dissemination of *C. neoformans*

Patrolling monocytes were recently shown to act as Trojan Horses and drive dissemination of *C. neoformans* into the CNS [19]. These studies were completed using an intravenous model of infection [23], and therefore the role of patrolling monocytes in extrapulmonary dissemination is unclear. Our data showed that PLX5622 caused a depletion of patrolling monocytes within the CNS (Fig 2), raising the possibility that the reduction in fungal burdens we observed in this tissue may in part be caused by a reduction in Trojan Horse-driven dissemination within patrolling monocytes. To explore this possibility, we first measured recruitment of patrolling monocytes to the CNS after intranasal infection. We found no significant increase in the number of patrolling monocytes in the brain or meninges at several time points post-intranasal infection (Fig 7A). Next, we used mice lacking patrolling monocytes to determine the role of these cells for anti-cryptococcal immunity following pulmonary infection. Nr4a1 (Nur77) is a transcription factor that controls the development of patrolling monocytes [30]. Mice lacking Nr4a1 have significantly reduced circulating patrolling monocytes [30], which we confirmed in our mice (Fig 7B). *Nr4a1*^-/-^ mice can therefore be used to assess the contribution of patrolling monocytes for the control of *C. neoformans* infection. We intranasally infected these animals with *C. neoformans* and measured fungal burdens in the brain and lungs at 21 days post-infection. We found no significant differences between *Nr4a1*^-/-^ mice and wild-type controls in control of lung or brain infection in either of these organs (Fig 7C). Taken together, we conclude that patrolling monocytes therefore play a redundant role in dissemination and control of *C. neoformans* infection following intranasal administration.

**Figure 7:**
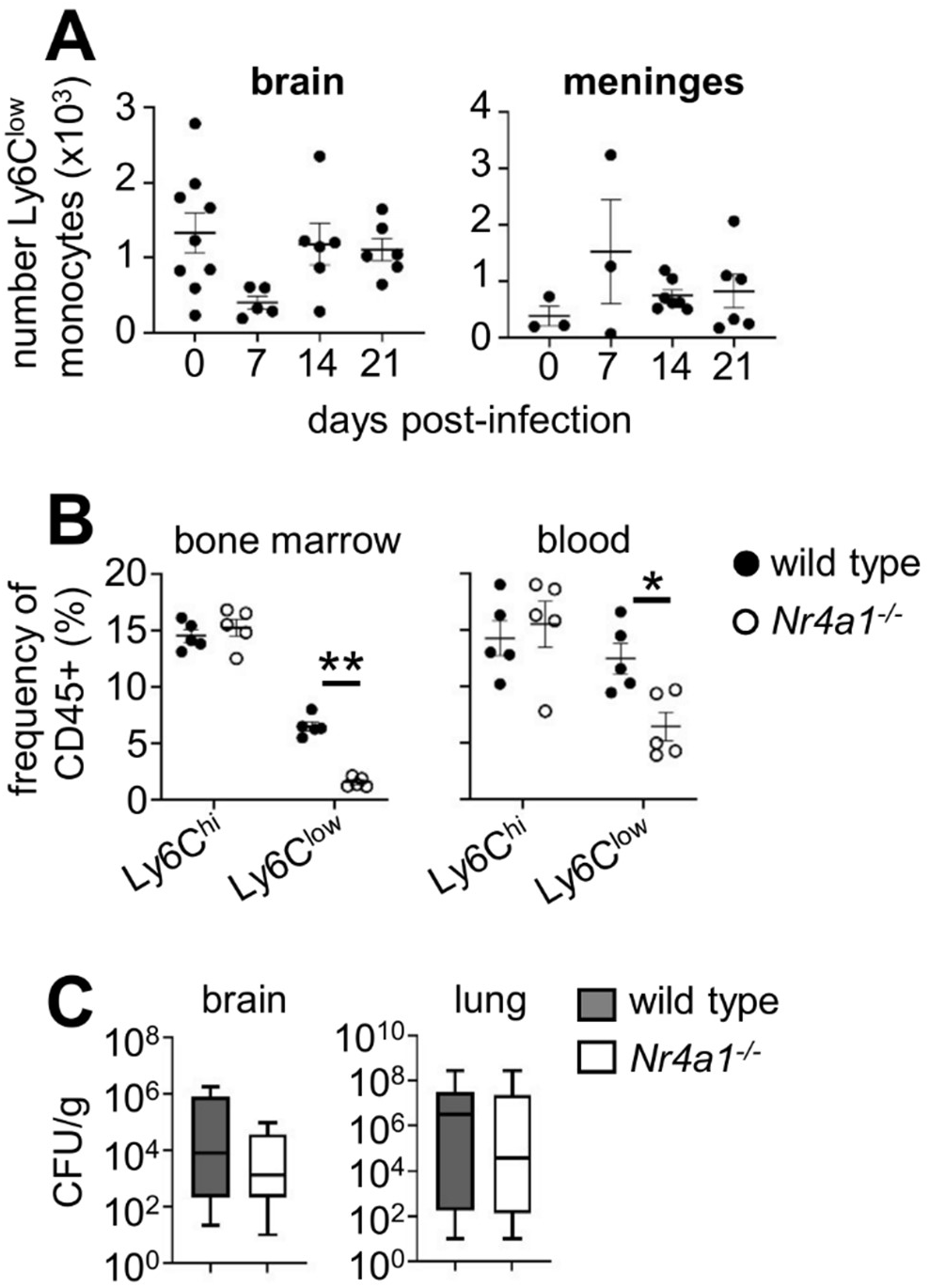
Patrolling monocytes play a redundant role in control and dissemination of pulmonary *C. neoformans* infections. (A) Total number of Ly6C^low^ patrolling monocytes in the brain and meninges of wild-type C57BL/6 mice at indicated time points post-intranasal infection. Each point represents a single animal. Data is pooled from 1 (meninges) or 2 (brain) independent experiments. (B) Frequency of Ly6C^hi^ and Ly6C^low^ monocytes in the bone marrow and peripheral blood of uninfected *Nr4a1*^-/-^ (n=5 mice) and wild-type controls (n=5 mice). Data from a single experiment and analysed by two-way ANOVA. *P<0.05, **P<0.01. (C) Fungal burdens in the brain and lung at day 21 post-infection in wild-type (n=19) and *Nr4a1*^-/-^ (n=20) mice. Data pooled from 3 independent experiments.

## Discussion

Intracellular survival within monocytes and macrophages is a critical part of the *C. neoformans* infection lifecycle. Understanding the factors that influence fungal intracellular residence has been a core goal for developing new treatment strategies for this infection, which are urgently needed because cryptococcal meningitis now accounts for 19% of all AIDS-related deaths globally [18]. In the current work, we found that depletion of CNS and lung resident macrophages with high dose PLX5622 treatment reduced *C. neoformans* infection. PLX5622 limited fungal infection within MHCII^hi^ interstitial lung macrophages, which expanded during infection following monocyte replenishment.

PLX5622 is primarily used to deplete CNS-resident macrophages, since microglia (and other types of CNS-resident macrophages) are highly dependent on the CSF1R for survival [31]. Rodents deficient in CSF1R lack these brain macrophages [3, 5], and a recent case report illustrated that human mutations in *CSF1R* resulted in complete loss of microglia and a consequent lethal disruption to brain development [4]. We recently demonstrated that targeted microglia depletion reduced *C. neoformans* infection in the CNS in a model of acute meningitis that bypassed pulmonary immunity [21]. In the current work, we used an intranasal infection model where yeast infection disseminates from the lung to other organs following breakdown of pulmonary barrier immunity. We found that loss of microglia in the CNS equated to reduced fungal brain infection, validating our earlier observations in the acute infection model [21]. However, further examination of PLX5622-treated mice revealed that fungal infection was also reduced in the lung, which may have exaggerated the reduced fungal brain burden by reducing the rate of extrapulmonary dissemination. Indeed, we found strong correlation between lung and brain burden in this model, in line with other reports [25]. Therefore, while microglia may support *C. neoformans* infection in the brain following intranasal infection, the therapeutic effect we observed with PLX5622 treatment may be enhanced by additional effects on dissemination patterns caused by PLX5622-mediated depletion of non-CNS myeloid cells.

Extrapulmonary dissemination of *C. neoformans* is further influenced by fungal morphotype. In the lung, *C. neoformans* forms large cells termed titan cells that are unable to be phagocytosed [32, 33]. As infection progresses, *C. neoformans* forms small cells that have greater propensity to disseminate to the brain, spleen and liver [34]. These small cells, called seed cells, were recently shown to have greater survival and accumulation within the CNS following pulmonary infection [34]. Whether PLX5622 affects *C. neoformans* morphotype in the lung and brain was not explored, although we did not observe any impact of PLX5622 on *C. neoformans* growth at a range of physiological concentrations that were previously measured in mice treated with high dose PLX5622 [24].

Dissemination of *C. neoformans* from the lung to other body sites is partially mediated by circulating monocytes [20]. Early experiments showed that infection of mice with *C. neoformans*-infected monocytes resulted in greater dissemination rates compared to mice infected with yeast alone [35], indicating that fungal-infected monocytes are better equipped to enter tissues and spread infection. Indeed, Ly6C^low^ patrolling monocytes were shown to carry *C. neoformans* into the CNS vasculature and contribute towards brain infection via a mechanism that required TNFR signalling [19]. That study used an intravenous model of infection, but the role of patrolling monocytes in promoting brain infection using other infection routes was not explored. We found that PLX5622 depleted patrolling monocytes specifically within the brain but not in the blood or other tissues. Other studies have also reported a depletion of monocytes within the CNS [10, 14]. Monocytes circulating in the blood do not appear to be affected by PLX5622 treatment, either in our study or in others [10, 16]. Taken together, these findings suggest that PLX5622 may not deplete Ly6C^low^ monocytes directly, but that the recruitment of these cells into the CNS may depend on the presence of microglia and CNS border macrophages. Since PLX5622-treated mice had reduced patrolling monocytes in the CNS, we tested whether this contributed towards the reduction in brain fungal burdens. However, we found that patrolling monocytes are not a significant driver of dissemination to the CNS following intranasal *C. neoformans* infection. Mice lacking Nr4a1 (which controls patrolling monocyte development [30]) were not better protected from infection, nor did we see any measurable accumulation of this monocyte subset in the CNS following intranasal infection. Therefore, the involvement of monocyte subsets in *C. neoformans* dissemination and infection may depend on route of infection and is an important consideration for future studies examining *C. neoformans* infection kinetics and the role myeloid cells play in shaping these patterns.

Inflammatory Ly6C^hi^ monocytes can also promote *C. neoformans* infections, particularly within the lung. Targeted depletion of CCR2-dependent inflammatory monocytes reduced fungal lung infection (following intratracheal instillation) and prevented dissemination to draining lymph nodes and had a moderate effect on brain dissemination [36]. The specific mechanism by which inflammatory monocytes enhance *C. neoformans* lung infection is unclear, but did not depend on adaptive immunity, eosinophils or arginase induction [36]. It is possible that by targeting monocytes, the replenishment and expansion of lung-resident macrophages was altered in these studies, which may have influenced lung fungal burdens. Indeed, total macrophages and dendritic cells were depleted in the lungs of CCR2-deficient mice [36]. We found that interstitial macrophages, particularly the MHCII^hi^ subset, significantly expanded during infection which was driven by monocytes since local proliferation of interstitial macrophages was decreased during infection. Our understanding of the ontogeny of interstitial macrophages is still evolving, but appears to require low levels of monocyte replenishment that increases with age and/or inflammation [37]. Indeed, the lung macrophage pool dynamically changes following infection as a result of monocyte infiltration, resulting in significant remodelling of the macrophage population that can influence responses to subsequent injuries [38].

We found that MHCII^hi^ interstitial macrophages were particularly susceptible to intracellular infection with *C. neoformans*, which correlated with a significant upregulation of arginase in this subset. This data independently validates recent findings from Madhani and colleagues who demonstrated that a *C. neoformans* secreted peptide termed CPL-1 was a strong inducer of arginase and M2 functional polarisation, particularly of lung interstitial macrophages [22]. We found that intracellular infection and arginase induction mapped to the MHCII^hi^ subset of interstitial macrophages. Functional interstitial macrophage subsets have now been described in multiple studies, which largely agree on the existence of MHCII^hi^ and MHCII^low^ interstitial macrophages [26, 27, 39]. MHCII^low^ interstitial macrophages co-express Lyve1 and CD206, primarily localise around blood vessels and appear to be critical regulators of inflammation and fibrosis since their absence exacerbates lung injury and leukocyte recruitment [39]. MHCII^hi^ interstitial macrophages appear to reside near nerves in the lung [26], although there is some variation in reported findings about the localisation and function of these cells [27] which may be due to a lack of consensus on the best way to identify and define interstitial macrophage subsets [37].

In our study, we found that treatment with high dose PLX5622 caused specific depletion of interstitial macrophages in the lung, particularly the MHCII^hi^ subset. Other studies have also reported potential effects of PLX5622 on lung macrophage populations although characterisation of specific subpopulations were not examined in those studies [11]. In contrast to our findings with PLX5622, mice deficient in CSF1R do not have significant alterations in lung macrophage populations [3, 5]. It is therefore likely that the PLX5622-mediated depletion we observed is due to dosage of the drug and a combination of direct effects on interstitial macrophages and indirect effects on replenishing monocytes. Experiments that have examined sensitivity of lung macrophages to CSF1 uptake have shown that interstitial macrophages more readily uptake CSF1 compared to alveolar macrophages, despite similar expression of the CSF1R between the two populations [2]. This work suggests that although alveolar macrophages express CSF1R, they may be less sensitive to CSF1R inhibitors and therefore resistant to depletion. In contrast, interstitial macrophages may have a greater dependency on CSF1R signalling to regulate their numbers and activation. Although the ontogeny of interstitial macrophages is somewhat contested, these cells do appear to require some input from monocytes for their maintenance and turnover. Monocyte turnover of interstitial macrophages has been shown to occur at low levels although conflicting studies have indicated that both Ly6C^hi^ and Ly6C^low^ subsets might play a role [27, 39]. In contrast, alveolar macrophages originate from foetal monocytes and do not require peripheral turnover, maintaining their numbers via self-renewal [37]. Whether PLX5622 interrupts monocyte replenishment of lung-resident interstitial macrophages is unclear, although our data argue that monocytes were unable to overcome PLX5622-mediated depletion even during infection, indicating that PLX5622 may interrupt monocyte-mediated replenishment. The factors influencing replenishment and contribution of monocytes to tissue-resident macrophage pools during infection is an important avenue to explore in future studies.

Protection against *C. neoformans* infections largely depends on the action of lymphocytes. IFNγ-producing CD4 T-cells are needed to prevent intracellular replication of the fungus within monocytes and macrophages, and activate fungal killing pathways such as induction of iNOS expression [17, 40]. B-cells produce antibodies which opsonise the fungus and enable efficient phagocytosis, which is otherwise hampered by the presence of the fungal capsule that shields against uptake by classic antifungal pattern recognition receptors such as Dectin-1 [40]. Studies that explored the off-target effects of PLX5622 have reported depletions of non-myeloid populations including T-cells and B-cells [11]. This has resulted in controversy in the field as to whether these effects are widespread or due to concentration and timing of PLX5622 treatment. We therefore extensively characterised lymphocyte numbers in multiple tissues before and during fungal infection in PLX5622-treated mice. However, we found no clear effects on the lymphoid population in PLX5622-treated mice, in any of the tissues examined. We therefore attribute the reduction in fungal burdens observed in PLX5622-treated mice to loss of myeloid cells and not any off-target effects on other cells that do not express CSF1R.

Our work indicates that targeting CSF1R-dependent cells may protect against *C. neoformans* infection. Susceptibility to *C. neoformans* infections in HIV-infected humans has been partly attributed to the CSF1-CSF1R axis [41]. A recent genome-wide association study found several polymorphisms upstream of the *CSF1* gene that associated with greater susceptibility to cryptococcal meningitis in an African-based cohort of HIV-infected patients [41]. Human PBMCs stimulated with heat-killed *C. neoformans* significantly upregulate *CSF1* expression, and pre-treatment of PBMCs with CSF1 boosted uptake and killing of the fungus [41]. Activation of monocytes/macrophages with CSF1 is therefore an important axis for protection. However, our data indicate that macrophages dependent on CSF1 for survival may instead promote infection, and targeting these cells with PLX5622 could be therapeutic. Indeed, depletion of CSF1R-dependent disease-associated macrophages is a strategy recently approved for treatment of tenosynovial giant cell tumours [42, 43]. PLX3397 (pexidartinib) removes CSF1R-dependent macrophages that contribute to the growth and inflammation within these rare tumours [44]. Future studies should take into consideration the potential beneficial and harmful effects of manipulating CSF1R signalling during *C. neoformans* infection, which will likely be influenced organ- and timing-specific effects.

In summary, our data reveal that high dose PLX5622 treatment has a therapeutic effect for *C. neoformans* infection by removing intracellular growth niches in the lung and brain. We show that PLX5622 may be a useful strategy to deplete lung interstitial macrophages, which we have used to demonstrate the importance of these cells in the pathogenesis of *C. neoformans* lung infection and extrapulmonary dissemination.

## Methods

### Mice

8-12 week old C57BL/6 mice (males and females) were housed in individually ventilated cages under specific pathogen free conditions at the Biomedical Services Unit at the University of Birmingham, and had access to standard chow and drinking water *ad libitum*. Animal studies were approved by the Animal Welfare and Ethical Review Board and UK Home Office under Project Licence PBE275C33. Wild-type mice were purchased from Charles River. *Cx3cr1-GFP-Ccr2-RFP* (strain number 032127) and *Nr4a1*^-/-^ (strain number 006187) mice were purchased from Jackson Laboratories and colonies bred and maintained at the University of Birmingham. Mice were euthanised by cervical dislocation at indicated analysis time-points, or when humane endpoints (e.g. 20% weight loss, hypothermia, meningitis) had been reached, whichever occurred earlier.

### PLX5622 treatment

PLX5622 (Plexxikon Inc. Berkley, CA) was formulated in AIN-76A rodent chow (Research Diets) at a concentration of 1200 mg/kg. Mice were provided with PLX6522 diet or AIN-76A control diet *ad libitum* for 1 week prior to infection, and continued throughout the infection study period.

### *C. neoformans* growth and mouse infections

*C. neoformans* strains used in this study were H99 and KN99α-mCherry [45]. Yeast was routinely grown in YPD broth (2% peptone [Fisher Scientific], 2% glucose [Fisher Scientific], and 1% yeast extract [Sigma]) at 30 °C for 24 hours at 200rpm. For infections, yeast cells were washed twice in sterile PBS, counted using haemocytometer, and 2×10^5^ yeast (in 20μL) pipetted onto the mouse nares under isoflurane anaesthesia. For analysis of organ fungal burdens, animals were euthanized and organs weighed, homogenized in PBS, and serially diluted before plating onto YPD agar supplemented with Penicillin/Streptomycin (Invitrogen). Colonies were counted after incubation at 37°C for 48 hours.

### *C. neoformans* growth curve in PLX5622

In some experiments, *C. neoformans* was grown in the presence of PLX5622 for 48 hours. *C. neoformans* was seeded into 96 well plates at 5000 yeast/well, in the presence or absence of PLX5622 (3-12μg/mL; see figure legends for final concentrations). Yeast growth was monitored by serial readings at OD600 using a FLUOstar Omega microplate reader. Data was averaged across 3 wells per growth condition.

### Isolation of brain leukocytes

Leukocytes were isolated from brain using previously described methods [46]. Briefly, brains were aseptically removed and stored in ice-cold FACS buffer (PBS + 0.5% BSA + 0.01% sodium azide) prior to smashing into a paste using a syringe plunger. The suspension was resuspended in 10mL 30% Percoll (GE Healthcare), and underlaid with 1.5mL of 70% Percoll. Gradients were centrifuged at 2450 rpm for 30 min at 4 °C with the brake off. Leukocytes at the interphase were collected and washed in FACS buffer prior to labelling with fluorophore-conjugated antibodies and flow cytometry analysis.

### Meninges digestion

The crown of the skull was aseptically removed and meninges were gently peeled from the underside and collected in 1.5 Eppendorf containing 400μL digest buffer (RPMI supplemented with 1mg/ml collagenase [Fisher scientific], 1mg/mL dispase [Sigma] and 40μg/mL DNAse [Sigma]). Meninges were digested in a 37 °C water bath for 30 minutes with intermittent shaking every 5-10 minutes. The meninges were filtered through a 70μm cell strainer and then centrifuged at 1500rpm for 5 min at 4°C. Cells were resuspended in 200μL FACS buffer and stored on ice prior to staining.

### Isolation of lung leukocytes

Lungs were aseptically removed and placed in 4mL digest buffer (RPMI, 10% FBS, 1% Pen/Strep, 1mg/mL collagenase, 1mg/mL dispase and 40μg/mL DNase). The lungs were incubated in a water bath for 40-60 minutes at 37 °C with intermittent shaking every 5-10 min. Lung tissue then was gently smashed using a syringe plunger and filtered through 100μm filter. Lung cells were then collected by centrifugation at 1400rpm for 7 minutes at 4°C. Red blood cells were lysed on ice using PharmLyse solution (BD). Samples were washed in 2mM EDTA/PBS, filtered through at 40μm filter into a fresh tube, and collected by spinning at 1400rpm for 7 minutes. Cells were resuspended in 200μL FACS buffer, stored on ice, and placed in a FACS tube ready for staining.

### Collection and preparation of peripheral blood

Mice were anesthetised with isoflurane and up to 300μL of peripheral blood obtained via cardiac puncture. Blood samples were missed 50μL of 100mM EDTA/PBS solution and kept on ice. Red blood cells were lysed on ice for 5 minutes using 5mL of PharmLyse solution (BD), with gentle inversion to mix after 2.5 minutes. 8mL of 2mM EDTA/PBS was added to each sample, gently inverted to mix, and cells collected by centrifugation at 1500rpm for 5 minutes at 4°C. The supernatant was discarded, and cells washed in FACS buffer prior to staining and analysis by flow cytometry.

### Isolation of spleen leukocytes

Spleens were aseptically collected and placed in ice-cold PBS. For analysis of myeloid populations, spleens were digested in digest buffer (as above) for 30 minutes at 37°C with intermittent shaking prior to smashing and washing. For analysis of lymphoid populations, spleens were directly smashed with a syringe plunger through a 70μm filter. Red blood cells were lysed on ice using PharmLyse solution (BD). Samples were washed in 2mM EDTA/PBS, filtered through at 40μm filter into a fresh tube, and collected by spinning at 1500rpm for 5 minutes.

### Collection and preparation of bone marrow leukocytes

Hind leg femurs were collected from mice and connective/muscle tissue removed. Ends of the femur were removed using scissors, and bone marrow flushed out by passing 2mM EDTA/PBS through the bone using a fine needle and syringe. Bone marrow was collected by centrifugation, and red blood cells lysed using PharmLyse as above. Cells were washed in FACS buffer prior to staining and analysis.

### Flow Cytometry

Isolated leukocytes were resuspended in PBS and stained with Live/Dead stain (Invitrogen) on ice as per manufacturer’s instructions. Fc receptors were blocked with anti-CD16/32 and staining with fluorophore-labelled antibodies was performed on ice for 15-60 minutes. Labelled samples were acquired immediately or fixed in 2% paraformaldehyde prior to acquisition. In some experiments, samples were fixed/permeabilised using the Foxp3 staining buffer set (eBioscience) prior to intracellular staining for arginase (A1exF5, eBioscience) and Ki67 (16A8, Biolegend). Anti-mouse antibodies used in this study were: CD45 (30-F11), CD11b (M1/70), CX3CR1 (SA011F11), MHC Class II (M5/114.15.2), F480 (BM8), Ly6G (1A8), Ly6C (HK1.4), SiglecF (S1007L), CD64 (X54-5/7.1), CD206 (C068C2), all from Biolegend and MerTK (D55MMER) from eBioscience. Samples were acquired on a BD LSR Fortessa equipped with BD FACSDiva software. Analysis was performed using FlowJo (v10.6.1, TreeStar).

### Measurement of cytokines in lung

Lungs were stored in 1mL sterile PBS supplemented with 0.05% Tween20 and protease inhibitor cocktail (Roche), then homogenised using a Stuart Handheld homogeniser (Cole Parmer). Homogenates were centrifuged twice to remove debris and the supernatants snap-frozen on dry ice and stored at −80°C. Detection of IL-13 and IL-4 in lung homogenates was determined using Duoset ELISA (R&D system) as per manufacturers instructions.

### Micro-computed tomography

In these experiments, 9 week old BALB/cAn females were infected with 500 CFU of luciferase-expressing *C. neoformans* KN99α [25]. Mice were anaesthetised with isoflurane (2.5% in 100% oxygen) and scanned in supine position using a dedicated small animal *in vivo* μCT scanner (Skyscan 1278, Bruker micro-CT, Kontich, Belgium). The following parameters were used: 50 kVp X-ray source voltage, 1 mm aluminium X-ray filter, 350 μA current, isotropic reconstructed voxel size of 50 μm and 150 ms exposure time per projection with 0.9° increments over a total of 220° angle, resulting in a total scanning time of ~3 min with measured radiation dose of 60-80 mGy [47]. μCT-images were acquired weekly, from week 0 to week 4 post-infection. Software provided by the manufacturer (NRecon, DataViewer, and CTan) was used to reconstruct, visualise, and process μCT data. For Hounsfield unit calibration, a phantom of air-filled 1.5 mL tube located within a water-filled 50 mL tube was scanned. For each animal at each timepoint, μCT-derived biomarkers of lung function and pathology (aerated -, non-aerated -, and total lung volume) were quantified by manually delineating a VOI on transverse images avoiding the heart and main blood vessels [48].

### Statistics

Statistical analyses were performed using GraphPad Prism 9.0 software. Details of individual tests are included in the figure legends. In general, data were tested for normal distribution by Kolmogorov-Smirnov normality test and analyzed accordingly by unpaired two-tailed t-test or Mann Whitney U-test. In cases where multiple data sets were analyzed, two-way ANOVA was used with Bonferroni correction. In all cases, *P* values <0.05 were considered significant.

## Acknowledgements

We would like to thank the technical staff at the Biomedical Services Unit (Birmingham) for their care and help with animal husbandry. We thank Dr Guillaume Desanti and support staff at the Flow Cytometry Unit at the University of Birmingham for their support with sorting and flow cytometry experiments. This work was funded by the Academy of Medical Sciences (SBF004_1008, awarded to RAD), Medical Research Council (MR/S024611, awarded to RAD and MR/T029765/1, awarded to GA), and Research Foundation Flanders (PhD studentship awarded to EV, 1SF2222N).

## Supplementary Figures

**Figure S1:**
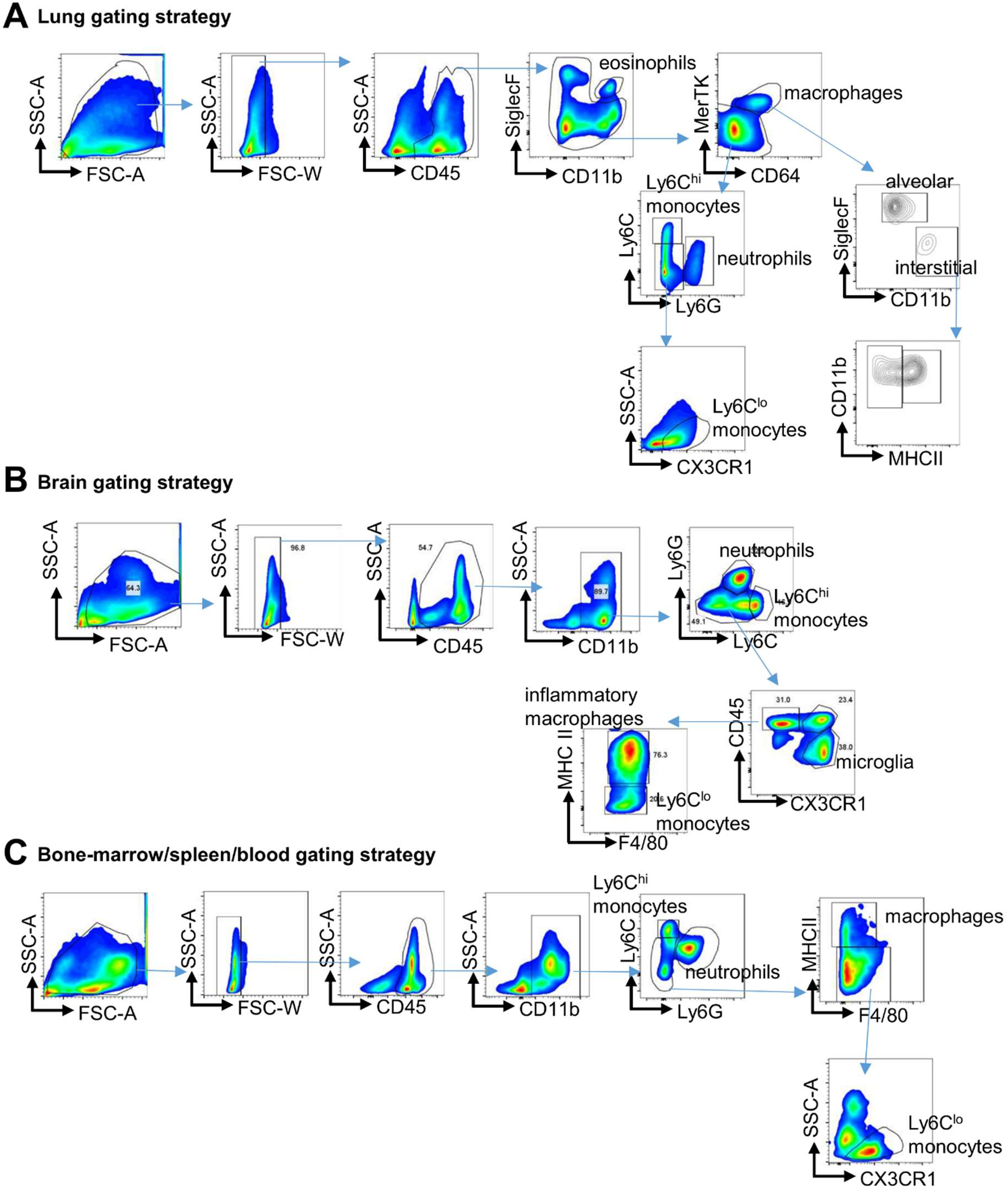
Gating strategies used in this study. Basic gating strategy used to define myeloid cell populations in the (A) lung, (B) brain and (C) spleen, bone marrow and blood.

**Figure S2:**
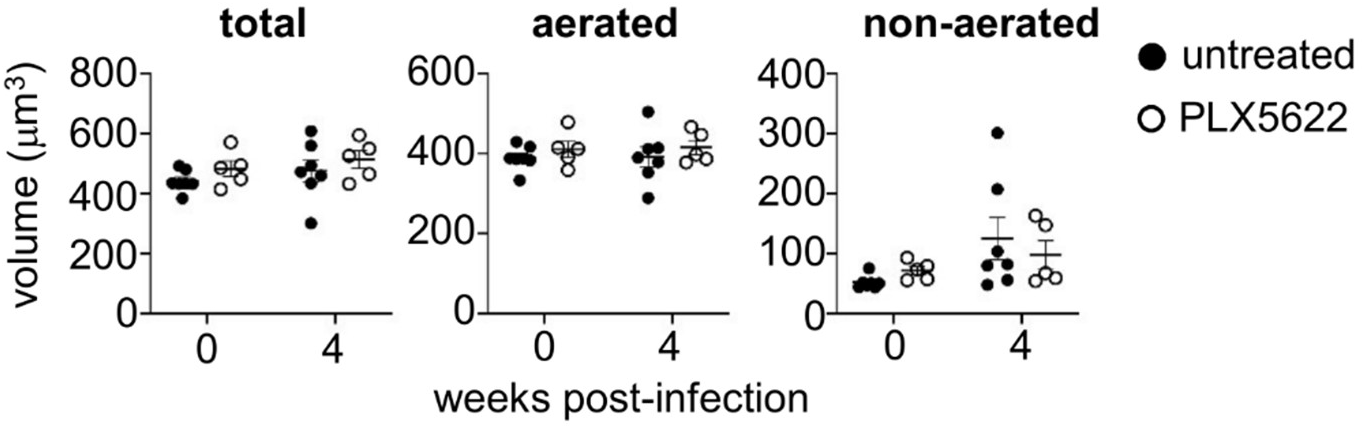
PLX5622 does not affect lung volume or aeration. Total lung volume, aerated lung volume and non-aerated lung (corresponding to lesion) volume were quantified from μCT images in mice that were untreated (n= 7 mice) or PLX5622 treated (n = 5 mice) at baseline (7 days post-diet, no infection) and 4 weeks post-infection. Data is pooled from 2 independent experiments.

**Figure S3:**
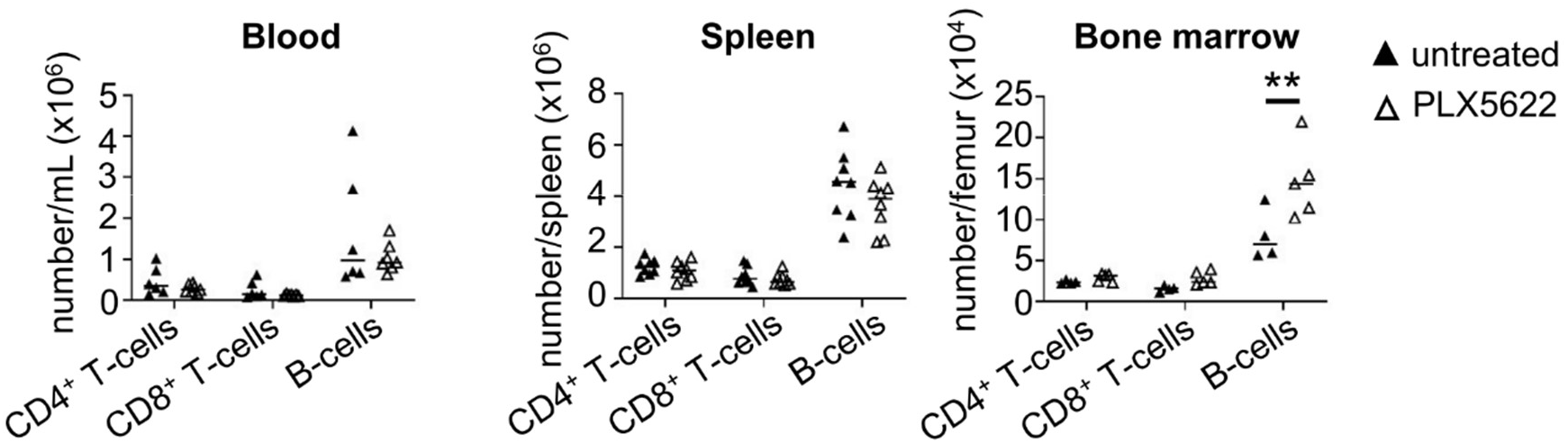
PLX5622 does not affect numbers of circulating lymphocytes. Total number of CD4+ T-cells, CD8+ T-cells and B-cells in the indicated organs of untreated (n=4-6 mice) and PLX5622-treated (n=5-8 mice) wild-type mice (all uninfected). Data is from 1 (bone marrow) or pooled from 2 independent experiments (blood, spleen).

**Figure S4:**
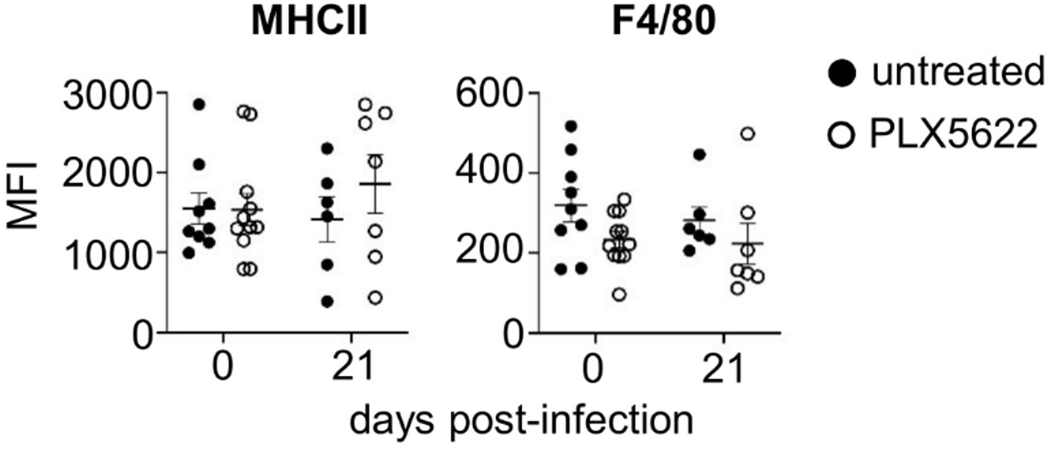
PLX5622 does not affect expression of MHCII or F4/80 by alveolar macrophages. Mean fluorescent intensity of MHCII and F4/80 by alveolar macrophages in the untreated (n = 6-9 mice) and PLX5622-treated (n = 7-10 mice) lung at the indicated time points relative to C. neoformans infection. Data pooled from 2-3 independent experiments.

**Figure S5:**
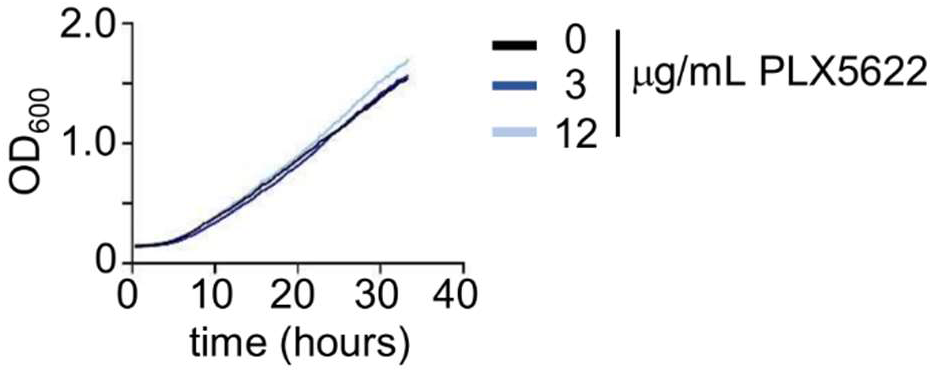
PLX5622 does not affect fungal growth directly. Growth curve of *C. neoformans* grown in the indicated concentrations of PLX5622. Data is representative of 2 independent experiments.

**Figure S6:**
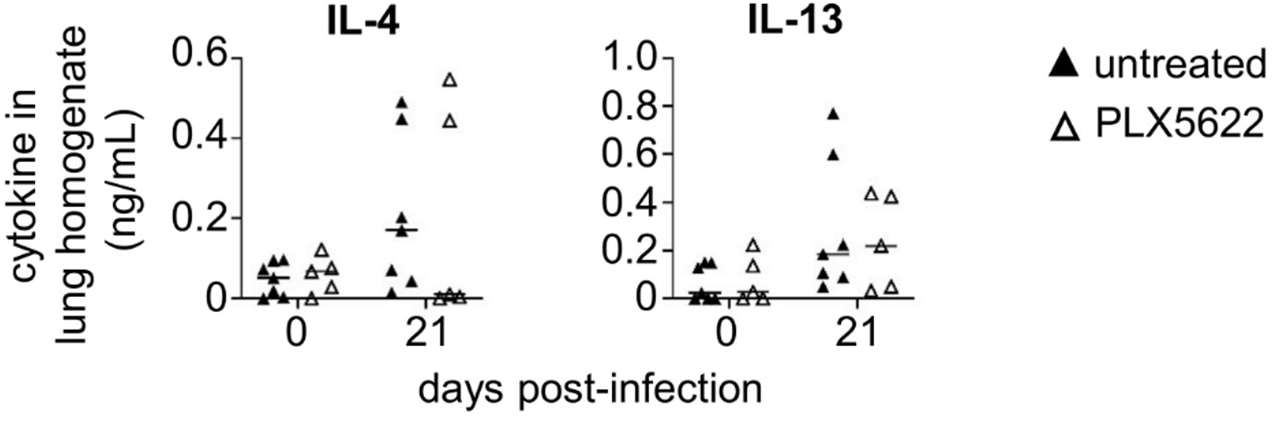
PLX5622 does not affect production of IL-4 and IL-13 within the lung. Concentration of IL-4 and IL-13 was measured in total lung homogenates of uninfected and infected mice, either untreated (n= 7 mice) or PLX5622-treated (n= 5 mice). Data pooled from 2 independent experiments.

**Figure S7:**
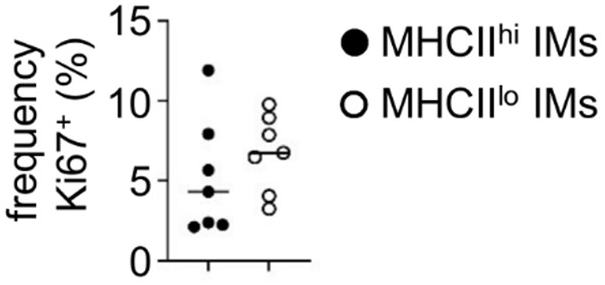
Interstitial macrophage subsets do not differ in Ki67 labelling. Frequency of Ki67+ interstitial lung macrophages that were either MHCII^hi^ or MHCII^lo^ IM = interstitial macrophage. Data pooled from 2 independent experiments (n= 7 mice).

**Figure S8:**
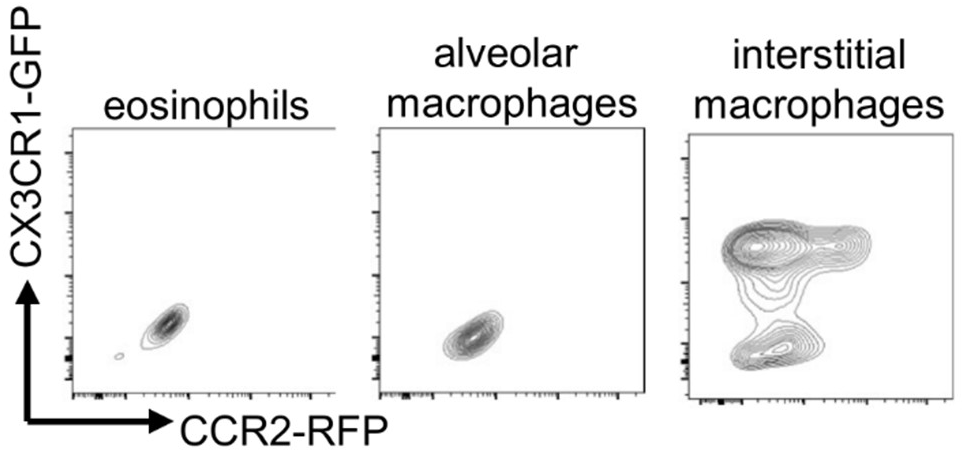
Background expression levels of CX3CR1 and CCR2 in the indicated cell types within the infected lung.

